# Loss of EED in the oocyte causes initial fetal growth restriction followed by placental hyperplasia and offspring overgrowth

**DOI:** 10.1101/2022.08.08.503175

**Authors:** Ruby Oberin, Sigrid Petautschnig, Tesha Tsai, Ellen G. Jarred, Zhipeng Qu, Neil A. Youngson, Heidi Bildsoe, Thi T. Truong, Dilini Fernando, Maarten van den Buuse, David K. Gardner, Natalie A. Sims, David L. Adelson, Patrick S. Western

**Affiliations:** Centre for Reproductive Health, Hudson Institute of Medical Research and Department of Molecular and Translational Science, Monash University, Clayton, Victoria, Australia; Department of Bioinformatics and Computational Genetics, School of Biological Sciences, University of Adelaide, Adelaide, South Australia, Australia; School of Biomedical Sciences, UNSW Sydney, Sydney, Australia; School of BioSciences, University of Melbourne, Parkville, Victoria, Australia; School of Psychology and Public Health, La Trobe University, Melbourne, Victoria, Australia; Bone Cell Biology and Disease Unit, St. Vincent’s Institute of Medical Research and Department of Medicine at St. Vincent’s Hospital, Fitzroy, Victoria, Australia.

## Abstract

Germline epigenetic programming, including genomic imprinting, substantially influences offspring development. Polycomb Repressive Complex 2 (PRC2) plays an important role in Histone 3 Lysine 27 trimethylation (H3K27me3)-dependent imprinting, loss of which leads to placental hyperplasia in mammalian offspring generated by somatic cell nuclear transfer (SCNT). In this study, we show that offspring from mouse oocytes lacking the Polycomb protein Embryonic Ectoderm Development (EED) were initially growth restricted, characterised by low blastocyst cell counts and substantial mid-gestational developmental delay. This initial developmental delay was followed by striking late-gestational placental hyperplasia, fetal catch-up growth and extended gestational length that culminated in offspring overgrowth. This involved remodelling of the placenta, including expansion of fetal and maternal tissues and conspicuous expansion of the glycogen enriched cell population in the junctional zone that was associated with a delay in parturition. Despite this remodelling and offspring catchup growth, fetal/placental weight ratio and fetal blood glucose levels were low indicating low placental efficiency. Genome-wide analyses identified extensive transcriptional dysregulation in affected placentas, including a range of imprinted and non-imprinted genes and increased expression of the H3K27me3-imprinted gene *Slc38a4,* which regulates transport of essential amino acids in the placenta. Our data provide an explanation for apparently opposing observations of growth restriction and overgrowth of offspring derived from *Eed-null* oocytes and demonstrate that PRC2-dependent programming in the oocyte regulates fetal and placental growth and developmental outcomes.

## Introduction

Epigenetic mechanisms orchestrate tissue development, body patterning and growth by regulating chromatin accessibility and expression of developmental genes in embryonic and extraembryonic tissues (1, 2). Histone 3 Lysine 27 trimethylation (H3K27me3) is a repressive epigenetic modification catalysed by Polycomb Repressive Complex 2 (PRC2), which is composed of core components, EED, EZH1/2 and SUZ12, all of which are essential for histone methyltransferase activity (3–6). Global deletion of any of the individual genes encoding these PRC2 subunits results in embryonic lethality in mice, while conditional deletions of *Eed, Ezh2* or *Suz12* result in altered patterning and organ function in a range of tissues (3, 7–13). In humans, *de novo* germline mutations in *EED, EZH2* or *SUZ12* result in Cohen Gibson, Weaver and Imagawa-Matsumoto syndromes, which all involve overgrowth, skeletal dysmorphologies and cognitive abnormalities in patients (14–20).

In mouse oocytes, PRC2 is required for non-canonical H3K27me3-imprinting, whereby H3K27me3 silences the maternal allele of several genes resulting in paternal allele-specific expression of the affected genes in pre-implantation embryos and extraembryonic tissues (21, 22). Oocyte-specific deletion of *Eed* using *Gdf9Cre* resulted in loss of H3K27me3-imprinting in preimplantation embryos and extraembryonic ectoderm with growth restriction and male-biased lethality in mid-gestation offspring (23). However, in contrast to growth restriction, another study that employed a similar strategy using *Zp3Cre* to delete *Eed* in oocytes found that postnatal offspring were overgrown (7). Moreover, somatic cell nuclear transfer (SCNT) embryos lack H3K27me3-imprinting, resulting in bi-allelic expression of several genes that has been linked to placental hyperplasia and fetal overgrowth (24–27).

In this study, we hypothesised that loss of EED in the oocyte establishes a developmental trajectory that involves initial fetal growth restriction followed by catch-up growth, explaining the overgrowth observed in early post-natal offspring. We demonstrate that while fetuses derived from *Eed* null oocytes were initially developmentally delayed and growth restricted, they underwent rapid catch-up growth that culminated in postnatal overgrowth. This complex growth curve occurred despite placental hyperplasia and reduced placental efficiency late in gestation. In addition, expression of H3K27me3-imprinted amino acid transporter *Slc38a4* was increased, gestational length was extended, contributing to the resolution of fetal growth restriction and ultimately resulting in postnatal overgrowth. Together, this work reveals an essential role for EED in the oocyte for regulating a complex program of fetal and placental growth in offspring, and a remarkable capacity for fetal growth restriction caused by loss of EED in oocytes to be resolved.

## Results

### Loss of maternal EED compromised mid-gestation survival in offspring

Previous studies demonstrated apparently contradictory findings in that deletion of *Eed* in oocytes resulted in early embryonic growth restriction (23) and early postnatal offspring overgrowth (7). To understand how these outcomes are realised, we produced offspring from pure C57BL6 females producing *Eed*-homozygous null (*hom*), *Eed* heterozygous (*het*) or *Eed* wild type (*wt*) oocytes and assessed their growth and development throughout the whole gestational period. Initially, *Eed^fl/fl^* or *Eed^fl/wt^* females were mated to males that were transgenic for *Zp3Cre* to generate females producing *Eed-wt, Eed-het* or *Eed-hom* oocytes (7). Mating of these females to pure C57BL6 *wt* males allowed us to generate isogenic wild type or heterozygous offspring as previously described (Fig. 1A) (7). Heterozygous offspring generated from *Eed* heterozygous oocytes (*HET-het* offspring) or from *Eed* homozygous null oocytes (*HET-hom* offspring) are isogenic. However, the *HET-hom* offspring were generated from oocytes that completely lacked functional EED and the *HET-het* offspring were generated from oocytes that had one functional copy of EED and maintained essentially normal gene repression (7(28). As they are isogenic and heterozygous, comparison of *HET-hom* with *HET-het* offspring allowed the identification of differences that resulted specifically from a loss of EED in oocytes in the absence of confounding genetic differences (Fig. 1A). In addition, we generated wild type offspring from *Eed-het* and *Eed-wt* oocytes to compare *WT-wt* and *WT-het* offspring with *HET-het* and *HET-hom* offspring, providing controls for differences generated as a result of *Eed* heterozygosity in offspring (Fig. 1A).

**Figure 1.**
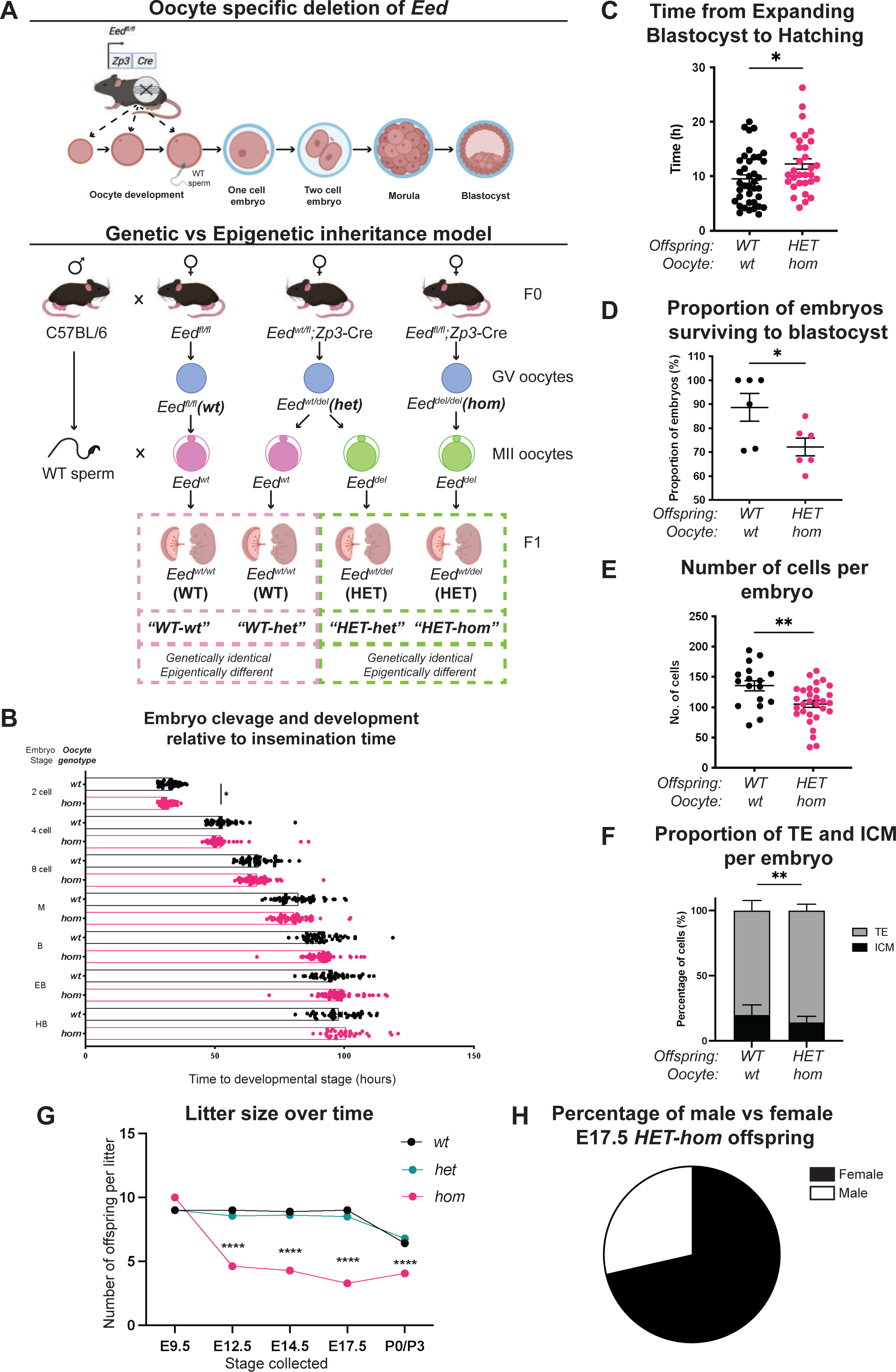
Maternal *Eed* deletion impacted pre-implantation development. (a) Model used to generate heterozygous isogenic offspring from oocytes that lacked or retained *Eed.* Females producing *Eed*-*wt*, *Eed*-*het* and *Eed*-*hom* oocytes were mated with wild type males to produce offspring from oocytes with *wt* or *het* EED-dependent programming or from oocytes that lack EED-dependent programming respectively. *Eed-wt* oocytes produce *WT* offspring (*WT-wt* control), *Eed-het* oocytes produce *WT* or *HET* offspring (*WT-het* and *HET-het*) from the same GV oocyte, and *Eed-hom* oocytes produce *HET* offspring (*HET*-*hom*). *HET-het* and *HET-hom* offspring are isogenic but were derived from oocytes that had different EED-dependent programming. **(b)** Cell cleavage and development times of embryos from *Eed-wt* or *Eed-hom* oocytes in *ex vivo* culture. Data is presented as time to reach 2-, 4-, 8-cell, morula (M), blastocyst (B), expanded blastocyst (EB) and hatched blastocyst (HB). *P<0.05, two-tailed student’s t test, N = 49 embryos from *Eed-wt* oocytes, 57 embryos from *Eed-hom* oocytes. **(c)** Time taken for expanded blastocysts to hatch. Data represents the time difference between hatched blastocyst and expanded blastocysts. *P<0.05, N = 38 embryos from *Eed-wt* oocytes, 31 embryos from *Eed-hom* oocytes. **(d)** Proportion of embryos from *Eed-wt* or *Eed-hom* oocytes surviving to blastocyst. *P<0.05, Data represent the proportion of surviving embryos from each female, N = 6 females. **(e)** Total number of cells per embryo. **P<0.005, N = 17 embryos from *Eed-wt* oocytes, 30 embryos from *Eed-hom* oocytes. **(f)** Proportion of TE and ICM cells per embryo. Data represents the mean proportion of cells allocated to TE versus ICM for embryos from *Eed-wt* or *Eed-hom* oocytes. **P<0.05, N = 17 embryos from *Eed-wt* oocytes, 28 embryos from *Eed-hom* oocytes. **(g)** Offspring litter size over time. ****P<0.0001. **(h)** Pie chart depicting the proportion of male and female *HET-hom* offspring at E17.5. N=32. For **(b-g)** a two-tailed student’s t test was used, and error bars represent mean ± SD.

Initially, we used automated time-lapse imaging of individual embryos derived from *Eed-wt* and *Eed-hom* oocytes to track their development from zygote to hatched blastocyst stages. Embryos from *Eed-hom* oocytes reached the 2-cell stage 1.08h earlier than embryos from *Eed-wt* oocytes (Fig. 1B), but blastocyst expansion and hatching took 2.67h longer in *HET-hom* compared to *WT-wt* embryos (Fig. 1C). All other developmental milestones to blastocyst stage were similar between genotypes (Fig. 1B). Although, viability to blastocyst stage was lower in embryos from females producing *Eed-hom* compared to *Eed-wt* oocytes (72.17% vs 88.67%, Fig. 1D), the proportion of embryos surviving to expanded and hatched blastocyst stages did not differ significantly (Supp Fig. 1A).

To determine whether the cell content of blastocysts was affected, we performed cell counts in whole blastocysts and differential staining of inner cell mass (ICM) and trophectoderm (TE) cells (29) (Supp Fig. 1B). This revealed that overall cell content of embryos from *Eed-hom* females was lower than in *Eed-wt* controls (Fig. 1E). The number of TE cells was not statistically altered, but the ICM contained fewer cells in *HET-hom* embryos compared to *WT-wt* controls (Supp Fig. 1C-D). Together, these data indicated that *HET-hom* embryos contained fewer cells overall than controls and that the ICM was smaller in embryos from oocytes that lacked EED compared to oocytes that maintained EED function (Fig. 1F). As previous reports indicated that maternal deletion of *Eed* did not compromise development to the blastocyst stage, or cause apoptosis in blastocysts (23), it is likely that lower proliferation explains the reduction we observed in blastocyst cell number, which affected the ICM rather than the TE.

To assess the effect of oocyte-specific *Eed* deletion on fetal survival and throughout pregnancy, we compared the number of live fetuses and litter size in offspring generated from *Eed-hom, Eed-het* and *Eed-wt* oocytes at multiple stages during gestation and after birth. Using *Zp3*Cre to delete *Eed* in oocytes, we found no difference between genotypes in the number of implantations of live embryos in four pregnancies from *Eed*-null oocytes analysed at E9.5 (Fig. 1G), indicating that *HET-hom* preimplantation embryos implanted at similar rates and progressed through gastrulation. However, consistent with previous findings (7, 23), the number of live fetuses at E12.5, E14.5, E17.5, and the number of live born pups were both significantly lower for *Eed-hom* mothers than for *Eed-het* and *Eed-wt* control females (Fig. 1G). In addition, 71.43% of live fetuses in E17.5 *Eed-hom* pregnancies were female, indicating male-biased lethality of *HET-hom* offspring (Fig. 1H), consistent with previous studies (23, 30).

### Loss of EED in oocytes results in offspring post-implantation developmental delay with catch-up growth and subsequent postnatal overgrowth

These observations demonstrated that the major period of fetal loss of *HET-hom* offspring was between E9.5 and E12.5 in our model or by E6.5 in Inoue, Chen (23), which indicates that either placental function or embryo survival was compromised in both models. To determine whether differences occurred in the fetus and/or placenta, we assessed the growth and development of surviving mid-late gestation offspring. Embryonic day (E)9.5 *HET-hom* offspring from multiple litters were small compared to *WT*-*wt* offspring (Fig. 2A). Similarly, E9.5 *HET*-*hom* offspring, including extraembryonic tissues, were significantly lighter than *HET-het, WT-het* and *WT-wt* controls (Supp Fig. 2A). Subsequently, compared to *HET-het, WT-het* and *WT-wt* controls, *HET-hom* embryos contained fewer tail somites at E12.5 and had reduced inter-digital tissue regression in foot plates at E14.5 (Fig. 2B, Supp Fig. 2B), demonstrating that *HET-hom* offspring were developmentally delayed.

**Figure 2.**
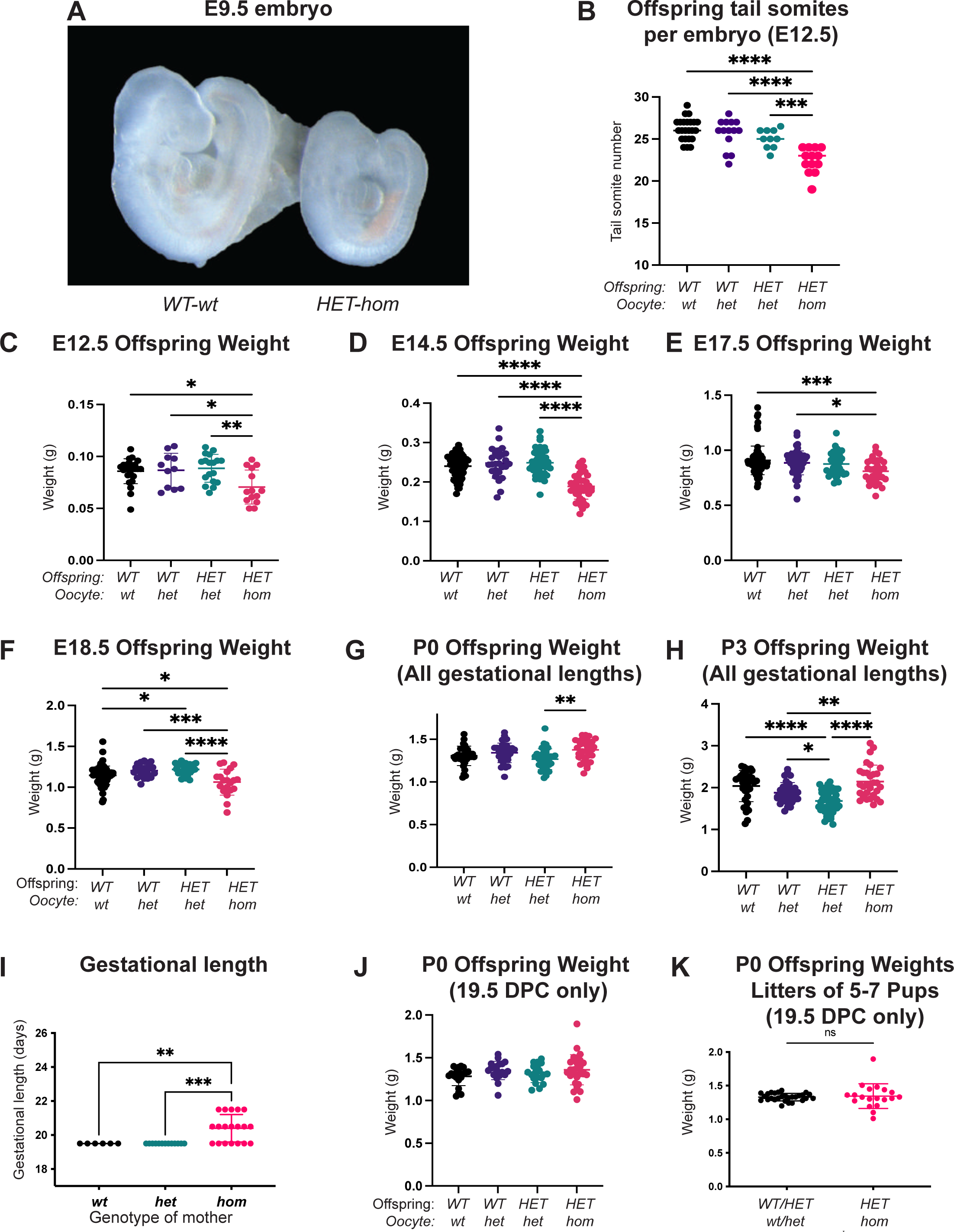
Loss of EED-dependent oocyte programming resulted in fetal loss and offspring developmental delay. (a) Representative wholemount images of E9.5 embryos. Images representative of 2-3 litters/genotype. **(b)** Number of tail somites in E12.5 *WT-wt, WT*-*het*, *HET-het* and *HET-hom* embryos. ****P<0.0001, ***P<0.0005, N=10-23/genotype. **(c-f)** Offspring body weight at E12.5, E14.5, E17.5 and E18.5. ****P<0.0001, ***P<0.0005, **P<0.005, N=11-75/genotype. **(g-h)** Offspring body weights at P0 and P3 from litters of all gestational lengths. ****P<0.0001, **P<0.005, *P<0.05, N=30-50/genotype**. (i)** Gestational length in litters from *Eed WT, HET* and *HOM* oocytes. **P<0.01, ***P<0.001. **(j)** Offspring body weight litters that were born on E19.5. N=19-28/genotype. **(k)** Offspring body weights of litters containing 5-7 pups and were born on E19.5. ns=non-significant difference, unpaired t test, N=19-28/group. **(b-j)** Statistically analysed using one-way ANOVA plus Tukey’s multiple comparisons.

This *in utero* developmental delay contrasted markedly with our previous observation that postnatal *HET-hom* offspring were overgrown at P2 (7). We therefore examined the temporal developmental trajectory of *HET-hom* offspring by measuring fetal weight at E12.5, E14.5 and E17.5 and pup weight on the day of birth (P0) and at P3. While weight of *HET-hom* offspring was significantly lower than *HET-het, WT-het* and *WT-wt* controls at E12.5, E14.5 and E17.5, by P0 and P3 *HET-hom* offspring were heavier than the isogenic *HET-het* controls (Fig. 2C-H). However, as we were collecting litters from time mated females, we also observed that gestational length was extended by 1 day in 7 pregnancies, or 2 days in 5 pregnancies of 19 pregnancies examined from *Eed-hom* oocytes, but gestational length was not extended in litters from *Eed-wt* and *Eed-het* oocyte controls (Fig. 2I). To determine the impact of this extended gestational time on offspring weight, we examined offspring weight of pups born on E19.5 days only. This revealed that offspring from *Eed-hom* oocytes were similar weight to controls (Fig 2J), indicating that *HET-hom* offspring growth had undergone catch-up growth and weight had been normalised by E19.5. Our previous report and data collected in this study demonstrated that litter size was also decreased from *Eed-hom* oocytes, raising the possibility that reduced *HET-hom* litter size may contribute to the normalisation of weight of pups at E19.5 (7). To address this, we compared pup weight from *Eed-hom* and control litters that contained 5-7 pups and were born on E19.5. This demonstrated that there was no difference in the weight of pups from *HET-hom* and control litters containing 5-7 pups (Fig 2K), also indicating that fetal growth restriction had been resolved by birth and that this catch-up occurred *in utero* and in a fashion that was independent of litter size. Together, while *HET-hom* offspring were initially developmentally delayed and under-weight compared to their isogenic heterozygous counterparts and wild type controls, they underwent rapid catch-up growth *in utero* and were overgrown by P3 (Supp Fig. 2C), consistent with our previous observations of overgrowth at P2 (7).

### Despite *HET*-*hom* offspring catch-up growth, oocyte-specific loss of EED results in late gestational placental hyperplasia and reduced placental efficiency

To understand whether the catch-up growth observed in *HET-hom* offspring might be related to changes in placental development, we collected and weighed E12.5, E14.5, E17.5 and E18.5 placentas from *HET-hom, HET-het, WT-het* and *WT-wt* offspring. At E12.5 placental weights were consistent across all genotypes, however at E14.5 *HET-hom* placentas were slightly, but significantly, heavier than *HET-het* controls (Fig. 3A-B). By E17.5 *HET-hom* placentas were 65% heavier than *HET-het* controls (Fig. 3C), and at E18.5 *HET-hom* placentas were 31.5% heavier than *HET-het* placentas (Fig. 3D), changes reflected in differing *HET-hom* and *HET-het* placental growth rates over time (Fig 3E-F). While *HET-hom* placenta growth rates were 0.79 times greater than *HET-het* placentas between E12.5 and E14.5, the growth rate of *HET-hom* placentas was 2.84 times greater than *HET-het* placentas between E14.5 and E17.5 (Fig 3E-F). Moreover, comparison of placental to fetal weights revealed that the growth of *HET-hom* placentas immediately preceded the catch-up growth of late-gestation *HET-hom* fetal offspring (Fig. 3E, G). While fetal growth rate was initially low between E12.5 and E14.5 and was similar between E14.5 and E17.5 in *HET-hom* offspring and *HET-het* controls, *HET-hom* offspring gained weight at 2.06 times the rate of *HET-het* controls between E18.5 and birth (E19.5) (Fig 3G).

**Figure 3.**
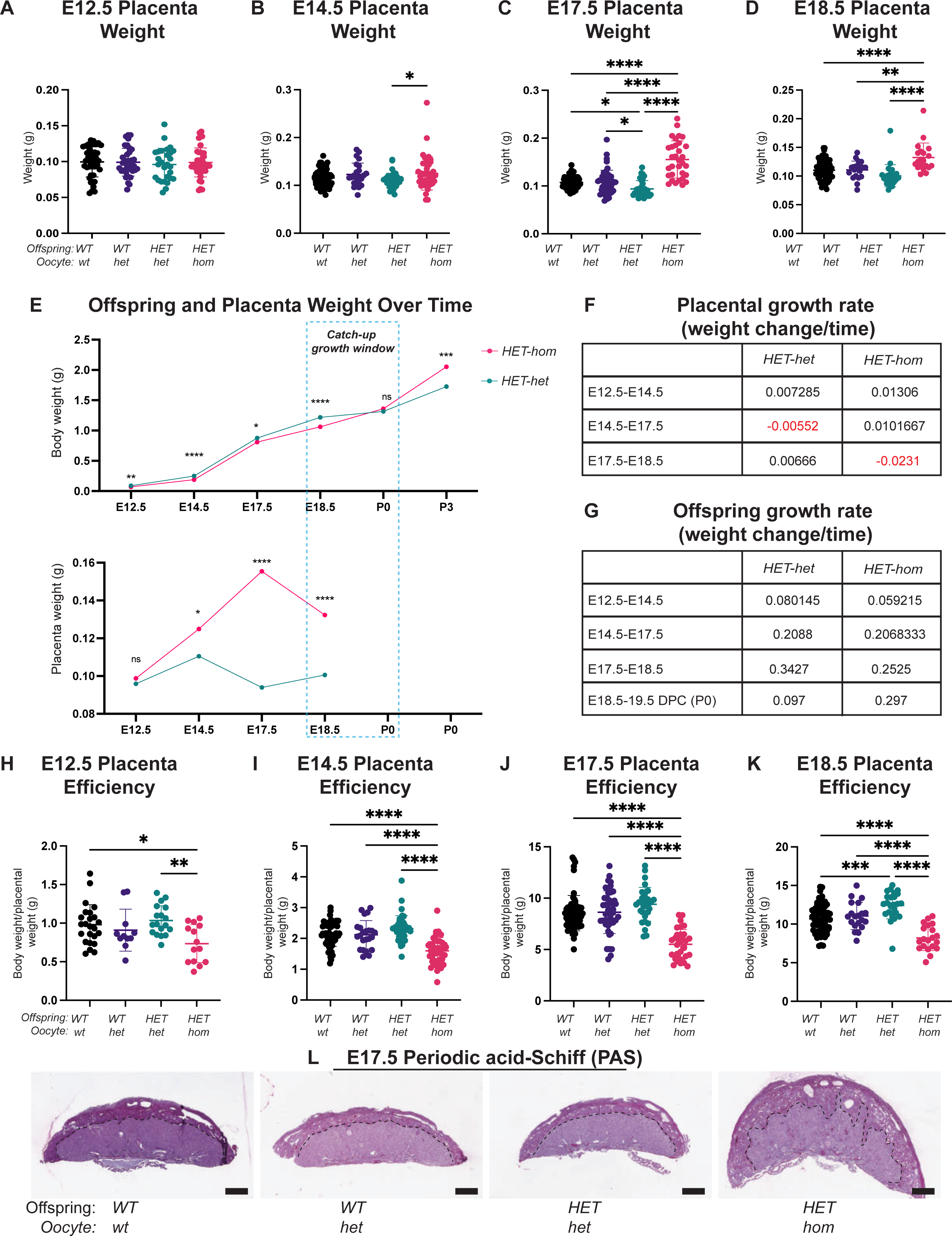
Placental differences were observed in heterozygous offspring from oocytes that lacked *Eed.* (a-d) Placental weight at E12.5, E14.5, E17.5 and E18.5, ****P<0.0001, **P<0.005, *P<0.05, N=20-66/genotype. **(e)** Isogenic heterozygous offspring and placenta average weight over time. ns=non-significant difference, *P<0.05, **P<0.005, ***P<0.0005, ****P<0.0001 unpaired t-test, N=28-56/genotype. **(f-g)** Tables depicting *HET-het* and *HET-hom* offspring and placenta growth rates across 3-4 time points throughout gestation. **(h-k)** Ratio of placental weight to offspring body weight at E12.5, E14.5, E17.5 and E18.5. ****P<0.0001, ***P<0.0005, **P<0.005, *P<0.05, N=11-66/genotype. **(l)** Placental cross-sections at E17.5 stained with PAS. Scale bars= 800μm. **(a-d, h-k)** statistically analysed using one-way ANOVA plus Tukey’s multiple comparisons.

To further investigate the relationship between placental function and offspring growth, we calculated the offspring body to placental weight ratio, which is indicative of placental efficiency (31). This indicated that placental efficiency was reduced at E12.5, E14.5, E17.5 and E18.5 in *HET*-*hom* offspring compared to all other genotypes (Fig. 3H-K). Histological sections at E17.5 confirmed that *HET-hom* placentas were obviously larger than *HET-het, WT-het* and *WT-wt* controls. This analysis also revealed that the junctional zone was expanded in *HET-hom* placentas, with abnormal projections of periodic acid-Schiff (PAS)-positive spongiotrophoblast cells into the labyrinth (Fig. 3J).

### Loss of EED in the oocyte results in altered developmental patterning of the placenta and extended gestational length

Given that PAS staining of E17.5 *HET-hom* placenta indicated substantial changes in placental development, we performed further analyses of *WT-wt*, *WT-het, HET-het* and *HET-hom* placentas at E14.5 and E17.5 using hematoxylin and eosin (H&E) staining. Consistent with the modest increase in weight of *HET-hom* compared to *HET-het* placentas at E14.5 (Fig 3B), quantification of the entire cross-sectional area in H&E-stained sections of E14.5 placentas demonstrated a 19% increase in *HET-hom* placental area compared to *HET-het* controls (Supp Fig. 3A & E). This difference was emphasised in overall cross-sectional area in E17.5 *HET-hom* placentas compared to all other genotypes, with a 55% increase in cross-sectional area compared to *HET-het* controls (Fig. 4A-B). To understand if this increase in placental size was due to changes in a specific layer of the placenta, we measured junctional zone, labyrinth and maternally-derived decidua layer areas. At E14.5, junctional zone area was significantly larger in *HET-hom* placentas compared to the *WT-wt* and *HET-het* controls, however there was no difference the *HET-hom* decidua or labyrinth areas compared to all other genotypes (Supp Fig. 3B-D). At E17.5, the overall cross-sectional area of all layers of *HET-hom* placentas was greater than that of *HET-het* and *WT-wt* controls (Supp Fig 3E-G). However, as a proportion of the total placental area, only the junctional zone was significantly greater for *HET-hom* placentas compared to other genotypes (Fig. 4C-E). Together, these data indicated that the whole *HET-hom* placenta, including the maternal side, was larger than controls. Moreover, this was associated with a disproportionate expansion of the fetally-derived junctional zone between E14.5 and E17.5.

**Figure 4.**
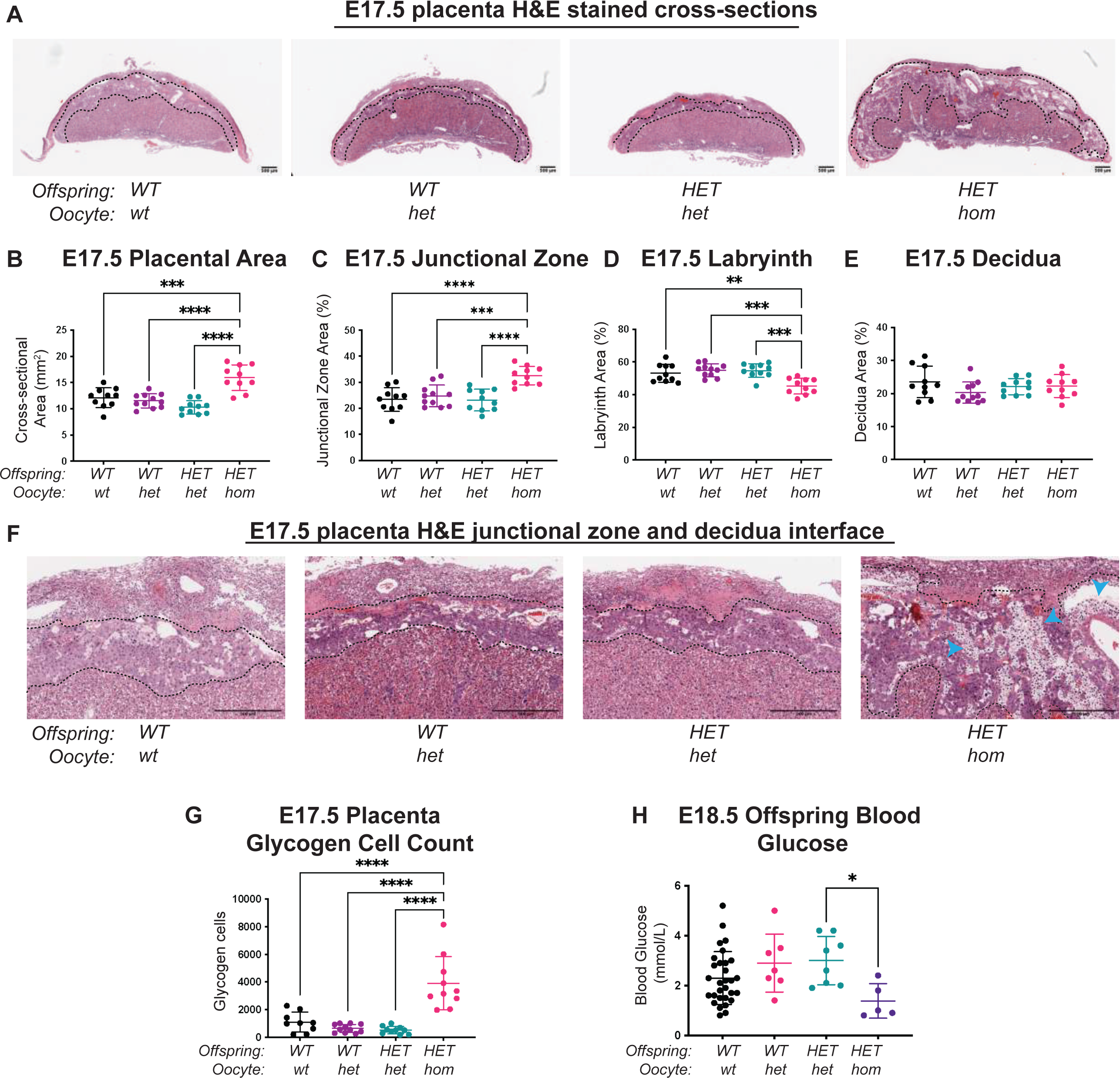
Complete deletion of *Eed* in the oocyte resulted in offspring altered placental morphology. (a) Placental cross-sections at E17.5 stained with H&E, black dotted line indicates the borders between labyrinth, junctional zone and decidual layers. Images are representative of four placentas from 10 biological replicates. Scale bars= 500μm. **(b)** Placental cross-sectional area at E17.5. ***P<0.0005, ****P<0.0001, N=10/genotype. **(c-e)** Area of E17.5 junctional zone, labyrinth and decidua. **P<0.005, ***P<0.001, ****P<0.0001, N=10/genotype. **(f)** Representative images of E17.5 junctional zone stained with H&E. Blue arrowheads, glycogen cells. Scale bars= 500μm. **(g)** Quantification of total number of glycogen cells in E17.5 placenta cross section. ****P<0.0001, N=10/genotype. **(h)** E18.5 offspring blood glucose concentration. *P<0.05, N=5-30/genotype. **(b-e, g-h)** statistically analysed using one-way ANOVA plus Tukey’s multiple comparisons.

Closer examination of H&E stained E17.5 placentas also showed that there were significantly more PAS-stained glycogen enriched cells counted in the entire *HET-hom* placenta midline sections compared to all other genotypes (Fig. 4F-G). To test whether these increased glycogen enriched cells provide an enhanced source of glucose to *HET-hom* offspring, we measured fetal blood glucose levels. However, consistent with decreased efficiency observed for *HET-hom* placentas (Fig 3H-K), *HET-hom* offspring had reduced blood glucose compared to *HET-het* controls, indicating that glucose levels are unlikely to contribute to *HET-hom* fetal catch-up growth (Fig 4H).

Placental glycogen cells are thought to play a role in inhibiting the production of hormones that promote parturition at term and glycogen cell number usually peaks in number at E16.5 in the mouse, and declines by E18.5, prior to delivery (32–34). It has been proposed that a higher number of placental glycogen cells towards the end of the pregnancy could delay the production of hormones that promote parturition (33–36). Consistent with this and the higher number of glycogen enriched cells in *HET-hom* placentas, gestational length was extended by 1 day or 2 days in *Eed-hom* pregnancies compared to the *Eed-wt* and *Eed-het* controls (Fig. 2I). However, in litters where gestational length was extended by one day from 19.5 days to days, 36.36% of the *HET-hom* offspring were found dead at birth, and in litters where gestational length was extended by two days to 21.5 days, all *HET-hom* offspring were found dead at birth. Together, junctional zone area and glycogen cell content was increased in all *HET-hom* placentas and gestational length was extended in almost 2/3 of all litters from oocytes lacking EED. Moreover, despite initial embryonic growth delay, reduced placental efficiency and reduced fetal glucose levels, *HET-hom* offspring underwent catch-up growth so that their weight was normalised by birth and was further enhanced in early postnatal life.

### Widespread gene dysregulation occurs in placentas from *Eed*-*hom* oocytes

As the developmental changes observed in the placenta indicated that transcriptional regulation may be substantially altered in *HET-hom* placentas we analysed male and female placental tissue from *HET-hom*, *HET-het* and *WT-wt* offspring using RNA-sequencing (RNA-seq). Surprisingly, principal component analysis of the male samples revealed that *HET-hom* placental transcriptomes were indistinct from placental transcriptomes in *HET-het* and *WT-wt* controls (Supp Fig. 4A), with only one sequence (CAAA01077340.1) differentially expressed between male *HET-hom* and *HET-het* placentas with an FDR<0.05 (false discovery rate) (Supp Fig. 4B). However, since we observed a male bias in fetal mortality in *HET-hom* offspring, the lack of extensive transcriptional differences between the male placental samples may have been influenced by more moderate effects in surviving males. We therefore focussed our analyses on female placentas only. Principal component analysis revealed that the transcriptome of female *HET-hom* placentas were distinct from their counterpart tissues in *HET-het* and *WT-wt* controls (Fig. 5A). Using an FDR<0.05, 2083 differentially expressed genes (DEGs; together referred to as *Eed* placental DEGs) were identified between *HET-hom* vs *HET-het* placentas (Fig. 5B, Supp Table 1), and only 4 genes (*Eed*, *Fibin*, *Hapln4* & *Dtx1*) were significantly different between *HET-het* and *WT-wt* placentas. This demonstrated that the vast majority of transcriptional dysregulation in the placenta was not due to heterozygosity of *Eed* but can be best explained by non-genetic effects caused by loss of EED specifically in the oocyte. Gene ontology analysis indicated that the *Eed* placental DEGs are involved in morphogenesis, system development and multicellular organism development (Fig. 5C), suggesting that the gene dysregulation observed in the *HET-hom* placenta influences a diverse range of systems and processes.

**Figure 5.**
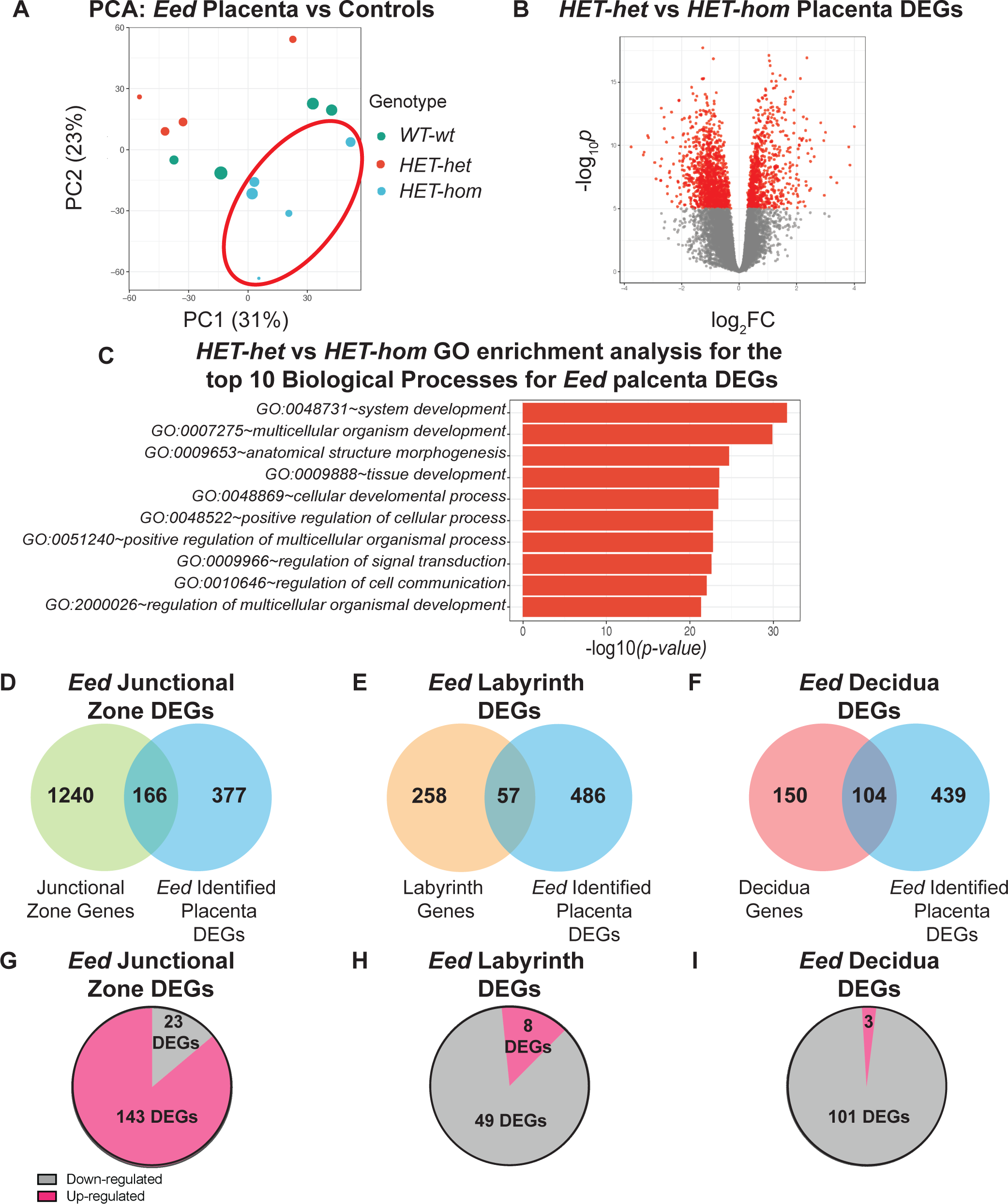
Loss of EED-dependent oocyte programming altered transcription in female *HET-hom* placenta. (a) Principal Component Analysis (PCA) for female *HET*-*hom*, *HET-het* and *WT-wt* placenta. N=4-5/genotype. **(b)** Differential gene expression analysis of isogenic *HET-het* vs *HET-hom* placenta represented by a volcano plot showing logFC against statistical significance. Genes with FDR-adjusted FDR<0.05 are coloured in red. Deletion of *Eed* in the oocyte resulted in 2083 DEGs in *HET-hom* placenta (*Eed* placental DEGs). **(c)** GO enrichment analysis of *HET-het* vs *HET-hom* DEGs representing the top 10 significantly different biological processes impacted (**d-f)** Venn diagrams showing comparisons of: **(d)** identified junctional zone genes vs *Eed* identified placenta DEGs, **(e)** identified labyrinth genes vs *Eed* identified placenta DEGs, **(f)** identified decidua genes vs *Eed* identified placenta DEGs. **(g-i)** pie graph depicting the distribution of up-regulated (pink) and down-regulated (grey) *Eed* DEGs identified to be expressed in **(g)** junctional zone, **(h)** labyrinth and **(i)** decidual layer of mouse placenta.

As sequencing was performed on the entire placenta, we compared the *Eed* placental DEG list with published single cell RNA-seq data from E14.5 C57BL/6 placentas as an indication of the placental cell types in which gene dysregulation occurred in our model (37). Although the E14.5 placental stage assessed by Han, Wang (37) was earlier than the E17.5 placental data collected in our analysis, the majority of mature placental cell types are established by E14.5 (38). We therefore considered the Han, Wang (37) data useful for predicting the spatial expression for our *Eed* placental DEGs. Of the 2083 *Eed* placental DEGs identified in the *Eed HET-hom* placentas, 543 genes were found in the E14.5 mouse placental data set (37). Cell-specific comparisons revealed that of these 543 genes, 166 *Eed* placental DEGs included genes preferentially expressed in the junctional zone and 57 *Eed* placental DEGs were preferentially transcribed in the labyrinth, which we defined as ‘*Eed* junctional zone DEGs’ and ‘*Eed* labyrinth DEGs’, respectively (Fig. 5D-E). In addition, 104 *Eed* placental DEGs included genes preferentially transcribed in the maternally-derived decidua, which we refer to as ‘*Eed* decidua DEGs’ (Fig. 5F). Together these data revealed that gene dysregulation in *HET-hom* placentas was not isolated to a single region, and that both maternal and fetal-derived cells were altered. Of the *Eed* junctional zone DEGs the majority (86.1%) were upregulated, whereas a large proportion of *Eed* labyrinth DEGs and *Eed* decidua DEGs were downregulated (86% and 97.1%, respectively; Fig. 5G-I). Together these data demonstrate a strong bias for up-regulation of genes expressed in the junctional zone layer of the *Eed* placenta. This reflects the significant expansion of the junctional zone as a proportion of the total placental size.

### *Eed* placental DEGs included both H3K27me3, classically imprinted genes and some X-linked genes

Given the loss of H3K27me3-dependent imprinting at specific genes leads to placental hyperplasia in Somatic Cell Nuclear Transfer (SCNT) embryos we investigated whether imprinted genes were dysregulated in the *Eed HET-hom* placenta. This revealed that of 325 known or proposed classically imprinted genes (39, 40), 46 were *Eed* placental DEGs (Fig. 6A, Supp Table 2). Of 76 putative non-canonical H3K27me3 imprinted genes identified by Inoue, Jiang (41), 16 were dysregulated in *HET-hom* placenta (Fig. 6A, Supp Table 1). These included, *Slc38a4*, *Sfmbt2*, *Gab1* and *Smoc1*, for which loss of H3K27me3-dependent imprinting results in placental hyperplasia and/or defective offspring development in SCNT-derived offspring (25–27). Moreover, *Slc38a4, Sfmbt2, Gab1 and Smoc1* were all up-regulated in *HET-hom* placenta, suggesting a loss of H3K27me3-imprinting and abnormal expression of the maternal allele of these genes (Fig. 6B). As a range of classically imprinted genes are also aberrantly expressed in placental cells of SCNT-derived pregnancies (27), we also determined the overlap of *Eed* placental DEGs with imprinted genes that were dysregulated in SCNT placentas. Of the 325 classically imprinted genes (39, 40), nine were commonly dysregulated (5 up-regulated and 4 down-regulated) in *HET-hom* placentas and SCNT derived placental cells. In addition, the H3K27me3-imprinted genes *Sfmbt2*, *Slc38a4, Smoc1* and *Gab1* were all upregulated in *Eed HET-hom* placentas, but only *Sfmbt2* dysregulated in placentas of SCNT-derived offspring.

**Figure 6.**
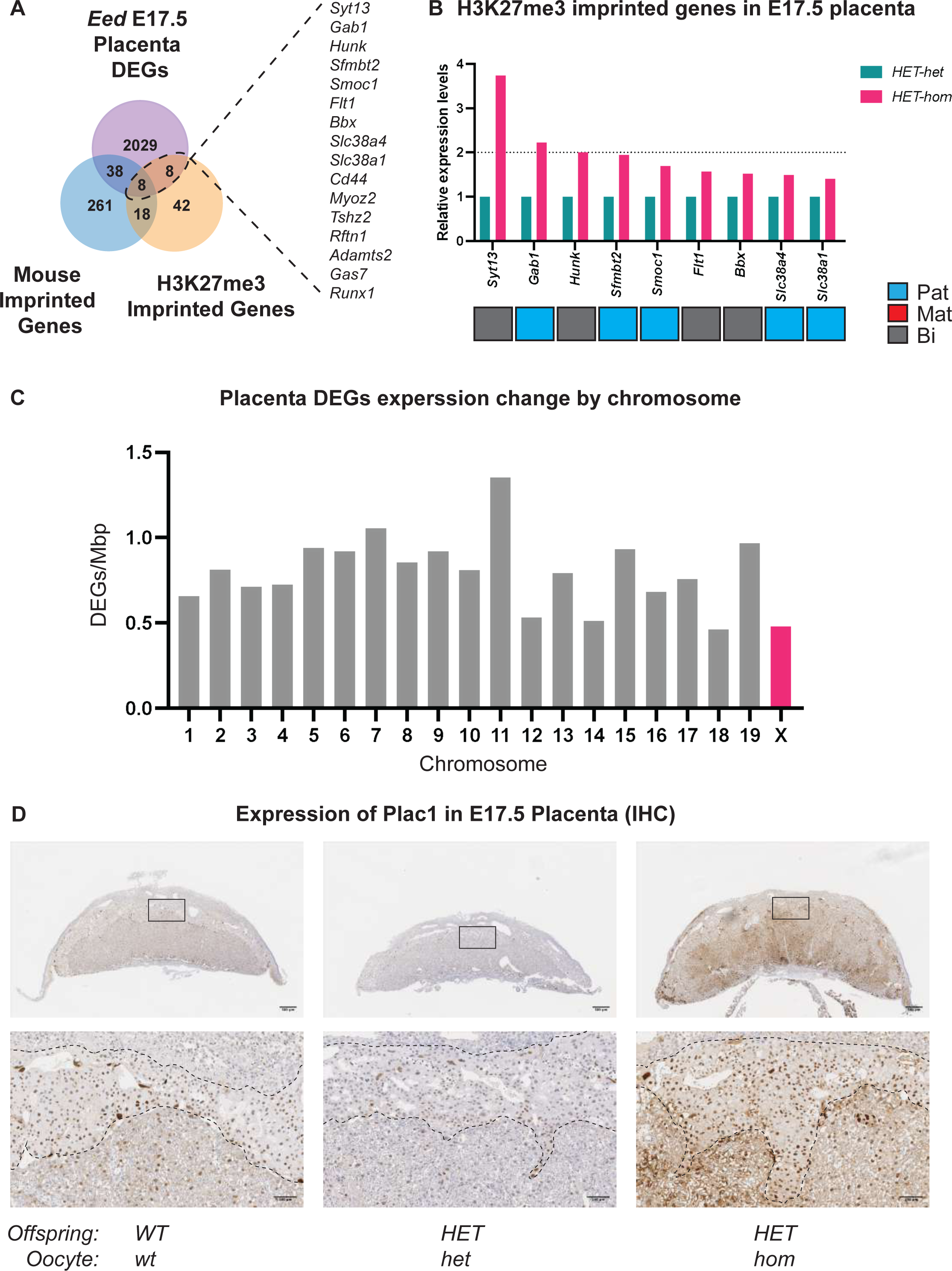
Loss of EED in the oocyte results in the de-repression of imprinted and X-linked genes. (a) Venn diagram showing comparisons of *Eed* placenta DEGs vs known mouse imprinted genes (39, 40) (bottom left) and H3K27me3 imprinted genes (22) (bottom right). **(b)** Relative expression levels of H3K7me3 imprinted genes that were upregulated in female E17.5 *HET-hom* placenta compared to female *HET-het* control (FDR<0.05). **(c)** *HET-hom* placenta DEGs expression change per chromosome. **(d)** Representative images of PLAC1 expression in E17.5 female placenta, n=4/genotype. Scale bars= 500μm (top), scale bars= 100μm (bottom).

As maternal EED is also required for regulating X-inactivation in pre-implantation embryos, and loss of EED in oocytes caused a male bias in fetal lethality in *Eed HET-hom* offspring (Fig. 1H) (23, 30), dysregulated X-inactivation may provide another explanation for the gene dysregulation detected in *Eed HET-hom* placenta. To determine if gene expression from the X-chromosome was preferentially impacted in *HET-hom* placentas, we examined the representation of the placental DEGs on all autosomes and the X-chromosome. Placental DEGs with increased expression occurred at a similar rate on the X-chromosome and autosomes, indicating that there was no preferential chromosome-wide de-repression of X-linked genes (Fig. 6C). However, we did note upregulation of X-linked genes *Plac1* and *Ldoc1,* and the imprinted genes *Ascl2* and *Peg3*, which are all known to regulate placental glycogen stores (33) (Supp Fig. 5A). All were increased in the junctional zone DEG list and have been associated with similar placental phenotypes to that observed in here *Eed Het-hom* placentas (Supp Fig. 5A) (42–47). In line with its increased transcription, PLAC1 protein was also increased in cells within the junctional zone (Fig. 6D).

Finally, we compared *HET-hom* growth patterns with available weight data reported for SCNT-derived offspring, in which H3K27me3 imprinting is disrupted (27). As for *Eed HET-hom* offspring, SCNT-derived offspring were underweight at E14.5, but had caught up by birth (Supp Fig. 6A-B), demonstrating that the growth profiles and placental phenotypes in these models are similar.

## Discussion

Independent studies have demonstrated that deletion of *Eed* in mouse oocytes using *Zp3Cre* or *Gdf9Cre* results in early growth restriction and offspring overgrowth outcomes in C57BL6 mice (7, 23). This study provides a cohesive explanation for these differing reports. We observed time-dependent and inter-related fetal and placental growth trajectories of offspring from *Eed-null* oocytes. Deletion of *Eed* in the oocyte initially led to embryonic and fetal developmental delay, followed by pronounced placental expansion and subsequent rapid catch-up and over-growth of the offspring realised in the early postnatal period.

Fetal growth restriction is commonly caused by placental insufficiency and is often investigated using surgical or nutritional interventions in animal models (48–51). In this study we provide a model that involves only the deletion of *Eed* in the oocyte and in which *HET-hom* offspring were compared to genetically identical *HET-het* controls. We detail fetal growth restriction immediately prior to placental hyperplasia. Despite lower placental efficiency, as measured by fetal/placental weight ratio, and low fetal glucose levels in *HET-hom* offspring, placental hyperplasia immediately preceded a period of catch-up growth. Similar placental hyperplasia and increased fetal weight outcomes were common between at least two models – the *Eed* oocyte deletion model described here and an SCNT model in which we found published data indicating similar growth restriction occurred prior to placental hyperplasia and subsequent fetal catchup growth was evident. Therefore, in both models, and potentially in others, the data support a model in which initial growth restriction is followed by placental hyperplasia and offspring catch-up growth, despite apparently lower placental efficiencies. In *Eed HET-hom* offspring, this was combined with extended gestational length that also contributed to increased birth weight. An imperative for offspring survival is for the delivery of sufficiently developed offspring. Considering this, an interesting conjecture may be that an unknown physiological mechanism that results in placental hyperplasia occurs alongside rapid intrauterine catch-up growth and prolonged gestation to rescue the pregnancy. While speculative, such a mechanism could involve epigenetic adaptation, perhaps H3K27me3-dependent or classical imprinting.

While this work has been in review, separate studies demonstrated that loss of *Eed* in oocytes results in placental hyperplasia due to loss of H3K27me3 imprinting at *Slc38a4* (52) and that late gestational placental hyperplasia, enhanced placental essential amino acid transport and increased fetal blood amino acid levels in SCNT-derived offspring is caused by loss of imprinting at *Slc38a4* (27). Consistent with these observations, we observed higher *Slc38a4* placental expression in *HET-hom* offspring, indicating that similar enhanced essential amino acid supply is likely to contribute to placental hyperplasia in *Eed HET-hom* mice. Therefore, while the fetal/placental ratio and fetal glucose levels were indicative of lower placental efficiency, increased amino acid transport across the placenta may explain the fetal catch-up growth in *HET-hom* mice.

The fetal growth restriction, placental hyperplasia and subsequent offspring overgrowth we observed in *Eed HET-hom* offspring was similar to that evident in SCNT-derived offspring. While both models involve loss of H3K27me3-dependent imprinting (23, 26, 27), the SCNT embryos are different to offspring from *Eed-*null oocytes because they were derived from oocytes containing functional maternal PRC2. This might indicate that altered H3K27me3-dependent epigenetic patterning of the oocyte genome is responsible for the early developmental delay in these offspring. However, the underlying cause could differ. SCNT embryos are derived by oocyte-driven reprogramming of a differentiated somatic nucleus, a process through which major epigenetic barriers must be overcome and which may compromise preimplantation development, particularly given that the embryos are transferred to recipient female mice (26, 53–55). Similarly, either loss of maternal PRC2 or H3K27me3 programming caused by *Eed* deletion in the oocyte could explain the reduced blastocyst cell content, and this may underlie early embryo and fetal growth restriction. One way to separate these issues may be to perform pronuclear transfer from *Eed-null* oocytes to wild type oocytes to rescue maternal PRC2 in offspring, and vice versa.

Comparison of isogenic *HET-hom* and *HET-het* offspring demonstrated that the phenotypic outcomes in this model originate from loss of EED in the oocyte and were largely unaffected by offspring genotype. While *HET-het* offspring maintained normal fetal growth, placental development, gestational length, and litter size, all these aspects of development were substantially modified in isogenic *HET-hom* controls. Other studies have reported that pre-implantation development proceeds normally in offspring from *Eed-null* oocytes, in that blastocysts formed at similar rates and were not affected by increased cell death (23). However, we found that blastocysts derived from *Eed-null* oocytes contained low cell numbers, and that this primarily affected the inner cell mass. Consistent with this, we observed substantial developmental delay in E9.5-E17.5 *HET-hom* offspring derived from *Eed-hom* oocytes, but this was resolved by birth and perinatal offspring were ultimately larger than controls.

There are at least two likely explanations for the early developmental delay observed in this model. EED supplied in the mature oocyte is rapidly localised to the maternal and paternal pronuclei within the first few hours following fertilisation (23, 28, 30, 56, 57). As the *HET-hom* offspring derived from *Eed-null* oocytes lacked maternal EED, it is possible that the lower cell numbers observed at the blastocyst stage resulted from low cell proliferation during pre-implantation development due to a lack of maternally-derived PRC2. This is consistent with the established role for PRC2 in driving cell division in stem cells and other cell types (58) and with a previous report that apoptosis was not increased in blastocysts from EED-null oocytes (23). Alternatively, the lower blastocyst cell counts we observed may have resulted from loss of EED-dependent epigenetic programming in growing oocytes. This programming occurs during a transient stage of PRC2 expression in primary-secondary stage follicles prior to the onset of DNA methylation (22, 28, 59).

We observed widespread morphological changes in *HET-hom* placental tissue. Placental expansion was observed in both fetally- and maternally-derived placental layers, indicating that while placental hyperplasia may be driven through the spongiotrophoblast cells in the junctional layer, the maternal side of the placenta also expands to accommodate greater placental function. Despite this, there was a disproportionate increase in size of the fetally-derived junctional zone compared to the maternally-derived decidual layer. Furthermore, closer examination of the enlarged junctional zone revealed an increase in glycogen enriched cells. This was unusual as glycogen-enriched trophoblasts peak in number at E16.5, but then decline by approximately 60% by E18.5 (33), an outcome reflected in the decreased placental weights we observed in E17.5 control offspring.

Given that fetal blood glucose levels were low, it seems unlikely that the higher number of glycogen cells in the E17.5 placentas of *HET-hom* offspring increase glycogen release to the embryo and drive rapid growth catch-up growth. In addition to facilitating increased glucose release and rapid fetal catchup growth, glycogen-producing trophoblasts are also considered to inhibit hormonal release from the placenta to the maternal blood supply to prepare the mother for parturition (33, 42). It may be that the increased number of glycogen cells remaining in late-gestation *HET-hom* placentas delay late gestational hormonal release and prolong pregnancy. Consistent with this, extended gestational length occurred in about two thirds of the pregnancies from *Eed-hom* oocytes. Moreover, while all pups were naturally born, the extended gestational length observed was also associated with high pup mortality in pregnancies extended by one day and mortality of all offspring in pregnancies extended by two days. Although the mechanisms underlying this extended gestation remain obscure, they appear to provide an extended widow for fetal/developmental catch-up in offspring, perhaps combining with *Slc38a4* and amino acid transport to increase the survival of live born pups.

In addition to the morphological alterations observed, we found extensive transcriptional dysregulation in *HET-hom* placentas, with 2083 genes differentially expressed compared to genetically identical *HET-het* placentas. This altered gene expression was not isolated to a single region of the *HET-hom* placenta and was observed in both fetal and maternal layers. While one might expect oocyte-specific loss of EED to affect only fetally-derived placental tissue, this ignores the possibility that the maternal tissue may expand or be re-modelled in response to placental hyperplasia in fetally-derived layers – indeed, without maternal re-modelling it seems likely that the placenta may not support fetal catch-up growth. On this basis, it is unsurprising that transcription was substantially altered in both fetally- and maternally-derived placental tissues.

Analyses of *HET-hom* placentas identified cellular and morphological changes in maternally- and fetally-derived tissues and placental DEGs that function in multiple systems. Recent work has demonstrated that loss of H3K27me3-dependent imprinting at *Slc38a4* or *Sfmbt2,* and to a lesser extent, *Gab1* or *Smoc1,* leads to placental hyperplasia in SCNT-derived offspring (25–27, 60). Consistent with this, we observed placental hyperplasia and increased expression of *Slc38a4, Sfmbt2, Gab1 and Smoc1* in placentas from *Eed-null* oocytes, suggesting a loss of H3K27me3-dependent imprinting at these genes. We also observed increased placental expression of the X-linked genes *Plac1*, *Wdr1* and *Ldoc1,* and immunohistochemistry confirmed that PLAC1 expression was increased in junctional zone cells in *HET-hom* placentas. However, deletion of *Plac1*, rather than increased expression, has been associated with a similar placental phenotype and male biased lethality to that observed in *HET-hom* offspring (61). Notwithstanding this difference, it seems likely that the fetal growth restriction, placental hyperplasia, and fetal catch-up growth in this model involves one or all of these genes, although additional work is required to understand the mechanisms involved and the interactions between the genes that contribute.

While we observed no difference in the number of surviving hatched or expanded blastocysts from *Eed-hom* mothers, there was a decrease in the blastocyst cell count, particularly in the ICM, and a marked decrease in *HET-hom* litter size. Consistent with this and with another study (23), we also observed smaller offspring at E9.5, developmental delay and loss of embryos by E12.5, and reduced litter size at term (7). While the reason for *HET-hom* offspring loss is yet to be identified, one possibility is that the significant reduction in blastocyst cell numbers and subsequent developmental delay is too profound to support survival during mid-gestation development in the most highly impacted offspring. This may involve impacts within the embryo itself, and/or contributions from a defective placenta. The placenta develops a capacity to exchange nutrients, other supportive metabolites, and waste between the maternal and fetal blood supply at E9.5, and increased embryo loss around this stage could indicate early-mid gestational placental insufficiency (62, 63). Embryonic lethality has also been previously linked to placental defects in knockout studies of imprinted and X-linked genes *Plac1*, *Ascl2* and *Peg10* (42-44, 46, 47, 64). As *Plac1* and *Ascl2* transcription was increased in the late-gestation *HET-hom* placenta, altered expression of imprinted genes such as *Plac1* or *Ascl2* may contribute to placental defects and fetal death in this model.

In summary, we demonstrate that loss of PRC2 function in the oocyte profoundly affects growth and developmental outcomes in both offspring and placenta via a mechanism that is independent of genetic background. This involves initial embryonic developmental delay and fetal growth restriction, followed by remarkable placental hyperplasia and fetal catch-up growth (Fig 7) associated with low placental efficiency and fetal blood glucose levels. Despite this, the fetal catch-up growth observed in this model may be explained by loss of imprinting for *Slc38a4* and increased placental essential amino acid transport. Importantly, rapid fetal growth restriction and fetal catch-up growth have been linked with negative health outcomes later in life, including metabolic conditions (65, 66). Moreover, given that mutations in the core PRC2 genes, EED, EZH2 and SUZ12 have all been associated with overgrowth and a range of co-morbidities in Cohen-Gibson, Weaver and Imagawa-Matsumoto syndrome patients (14, 15, 17, 19, 20) and we have observed similarities in outcomes of mouse offspring lacking EED in oocytes (7) this work also has implications for understanding these rare conditions. Together, this research reveals that altered PRC2-dependent programming in the oocyte elicits a complex intrauterine environment for the fetus, which has the potential to mediate impacts on offspring health and disease that may persist into adulthood.

**Figure 7.**
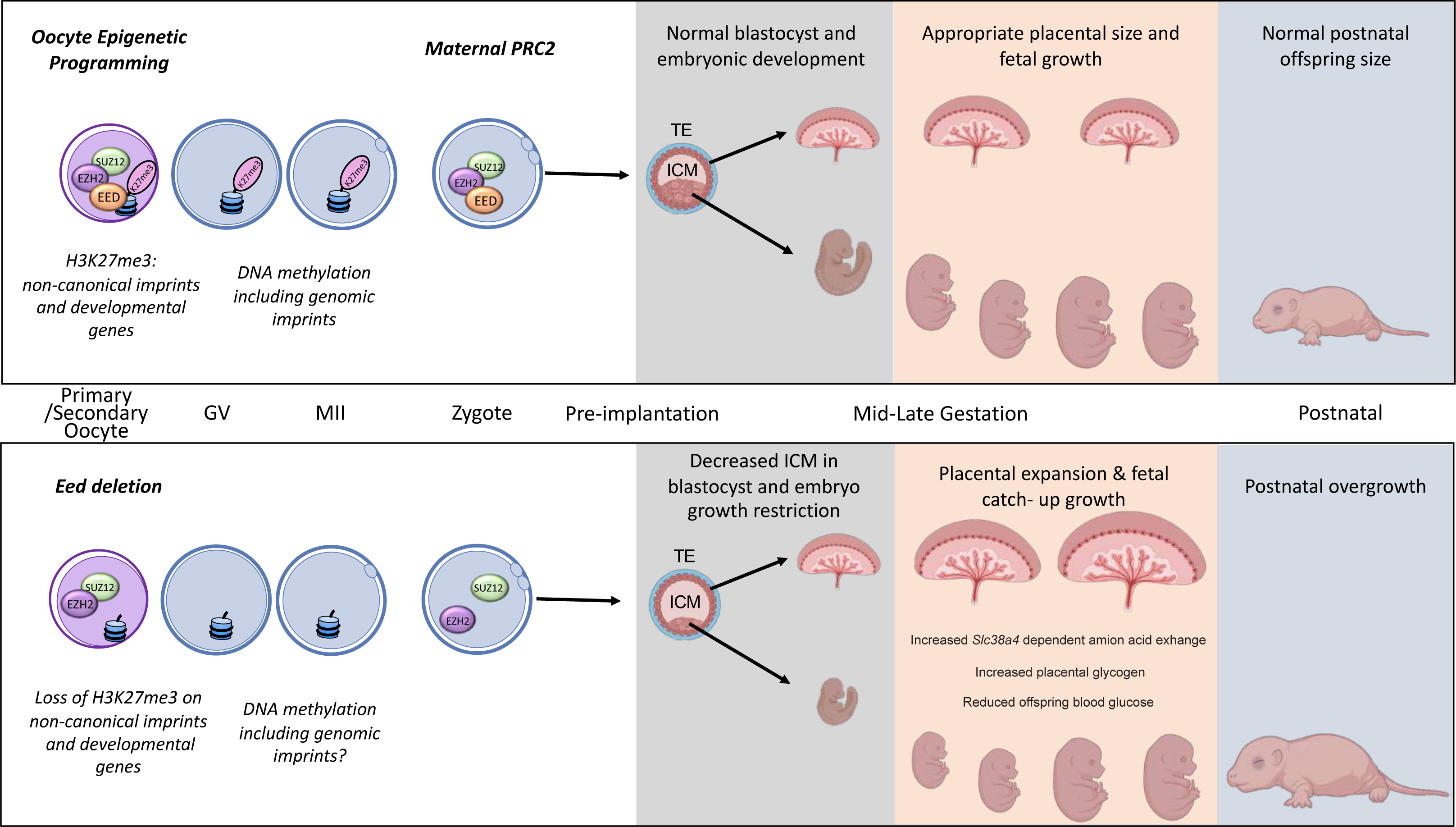
Summary of the placental and offspring growth response due to EED loss in the oocyte. During oocyte growth all three subunits of PRC2 are present at the primary to secondary stages, which is important for the silencing of developmental genes and H3K27me3-dependent imprinting (28). Deletion of *Eed* in oocyte results in loss of maternal H3K27me3 and PRC2, early growth restriction, followed by placental hyperplasia and late gestation fetal catch-up growth, outcomes consistent with loss of H3K27me3-dependent imprinting observed in SCNT offspring (25–27). Placentas generated from oocytes lacking EED have expanded glycogen enriched cells in the junctional zone and significant gene dysregulation in the placenta. Despite placental hyperplasia and reduced placental efficiency late in gestation, offspring catch-up growth observed in this model may be explained by loss of imprinting for *Slc38a4*, increased placental amino acid transport and extended gestational length, explaining why these offspring are overgrown immediately after birth (7).

## Methods

### Mouse strains, animal care and ethics

Mice were housed using a 12h light-dark cycle at Monash Medical Centre Animal Facility, as previously reported (7). Room temperature was maintained at 21-23°C with controlled humidity, and food and water were provided *ad libitum*. All animal work was undertaken in accordance with Monash University Animal Ethics Committee (AEC) approvals. Mice were obtained from the following sources: *Zp3Cre* mice (C57BL/6-Tg 93knw/J; Jackson Labs line 003651, constructed and shared by Prof Barbara Knowles (67), *Eed* floxed mice (*Eed*^fl/fl^) (B6; 129S1-*Eed*tm1Sho/J; Jackson Labs line 0022727; constructed and shared by Prof Stuart Orkin (68). The *Eed* line was backcrossed to a pure C57BL6/J and shared with us by Associate Professor Rhys Allen and Professor Marnie Blewitt, Walter and Eliza Hall Institute for Medical Research, Melbourne.

### Genotyping

Genotyping was performed by Transnetyx (Cordova, TN) using real-time PCR assays (details available upon request) designed for each gene as described previously (7).

### Collection and culture of pre-implantation embryos

Eight to twelve-week-old female mice were superovulated and mated to C57BL/6 males for one night. Zygotes were collected in handling media (G-MOPS PLUS, Vitrolife) at 37°C (29, 69) denuded of cumulus cells with G-MOPS PLUS containing hyaluronidase. All embryos were washed in G-MOPS PLUS and embryo development kinetics was assessed using the EmbryoScope™ (Vitrolife) time-lapse imaging system. Embryos were cultured individually in 25μl of medium, with time-lapse images generated at 15-minute (min) intervals throughout the culture period.

### Cell allocation in blastocysts

Following EmbryoScope™ culture, differential staining was performed (70) in hatched blastocysts using propidium iodide to label TE nuclei, while leaving the ICM unlabelled. After fixation, embryos were treated with bisbenzimide to stain ICM and TE, whole-mounted in glycerol and imaged using an inverted fluorescence microscope (Nikon Eclipse TS100). Nuclei were counted using ImageJ.

### Collection of post-implantation embryos, placenta and postnatal offspring

Mice were time mated for two-four nights, with females plug checked daily. Positive plugs were noted as day E0.5. Pregnant females were euthanised and embryos collected at E9.5, E12.5 E14.5 and E17.5. E9.5 whole sacs were weighed, fixed in 4% PFA for 72h at 4°C degrees processed, paraffin embedded, sectioned at 5µm. Whole-mount images of E9.5 embryos were taken using a LEICA M80 light microscope with LEICA MC170 HD camera attachment. E12.5, E14.5 and E17.5 embryos and placentas were isolated and weighed. Placentas were then bisected, and half was fixed in 4% PFA for 72h at 4°C degrees, processed and paraffin embedded with the cut side of the placenta facing the front of the block. The other half of the placenta was rinsed in PBS, snap-frozen on dry ice and stored at -80°C for RNA analysis. P0 and P3 offspring were weighed and euthanized by decapitation.

### Placental Histology

Paraffin embedded placentas were sectioned at 5µm using a Leica microtome and sections transferred to Superfrost plus slides (Thermo-Fisher, Braunschweig, Germany). Periodic antigen-shiff (PAS) and hematoxylin and eosin (H&E) staining was performed on placental sections by the Monash Histology Platform (MHTP node). Stained slides were scanned using Aperio slide scanner by the Monash Histology Platform and analysis was conducted using QuPath v0.2.3 (71). Histological analysis was conducted on one section located in the midline of each placenta. Junctional zone, labyrinth and decidual area was calculated in E14.5 and E17.5 H&E stained placentas. Glycogen cell counts were also conducted on the entire midline section of the H&E stained E14.5 and E17.5 placentas using QuPath v0.2.3 (71) . Investigators were blinded for sample genotypes throughout quantitative scoring of placental samples analysed.

### Immunohistochemistry

Slides with 5µm thick placental sections were baked at 60°C for 20 mins. Tissue sections were dewaxed in three changes of xylene and rehydrated in three changes of ethanol then rinsed in distilled water. Antigen retrieval was performed in DAKO PT Link in a DAKO Target Retrieval (Low pH) Solution (DAKO, Cat# S1699) at 98°C for 30 min. Slides were then washed in DAKO EnVision Flex Wash Buffer (Cat# K8000) for 5 minutes. IHC was then performed on a DAKO Autostainer Plus in the following steps. Sections were washed once in EnVision Flex Wash Buffer following each subsequent step. Peroxidase Blocking Solution (DAKO, Cat# S2023) was applied for 10 minutes and non-specific binding was prevented with AffiniPure Fab Fragment Goat Anti-Mouse IgG for 1 hour. Mouse anti-PLAC1 (G-1) (Santa Cruz, Cat# sc-365919) primary antibody was applied at an appropriate concentration for 1 hour. EnVision System-HRP Labelled Polymer Anti-Mouse (DAKO, Cat# K4001) was applied for 1 hour. Immunostaining was visualised using DAKO Liquid DAB+ Substrate Chromogen System (Cat# K3468). A counterstain DAKO Automation Haematoxylin Staining Reagent was then applied for 10 minutes. Slides were removed from the Autostainer, transferred to a slide staining rack and rinsed in distilled water. In a fume-hood, slides were then washed in Scott’s Tap water and distilled water. Finally, slides were dehydrated in three changes of 100% Ethanol, cleared in three changes of Xylene and mounted in DPX. Slides were either scanned using a VS120 Slidescanner (Olympus).

### Placental RNA-sequencing

Placental isolation is described above, and placental RNA was extracted from 4-5 samples/genotype using NucleoSpin RNA Plus columns. RNA quality was assessed on an Agilent Bioanalyser and samples with RIN >7.5 used for library preparation and sequencing on the BGI Genomics platform (BGI Genomics, Hong Kong).

### RNA-sequencing data analyses

Adaptor and low-quality sequences in raw sequencing reads were trimmed using Trimmomatic (72) (v0.39) with the following parameters: LEADING:3 TRAILING:3 SLIDINGWINDOW:4:15 MINLEN:20. Clean reads were mapped to the mouse reference genome (GRCm38) using STAR (v2.7.5c) with the following settings: outFilterMismatchNoverLmax 0.03 --alignIntronMax 10000. Raw counts for mouse reference genes (ensembl-release-101) were calculated using STAR (v2.7.5c) with parameter “-- quantMode GeneCounts” simultaneously when doing the genome mapping. Differential gene expression analysis was carried out using the R package “limma” (73) with “treat” function and parameter “lfc=log(1.1)”. Statistically significantly differentially expressed genes were identified using “FDR < 0.05”. Gene Ontology (GO) enrichment analysis for significantly differentially expressed genes was carried out using The Database for Annotation, Visualisation and Integrated Discovery (DAVID) with following settings: GO term level 3, minimum gene count 5, and FDR < 0.05 (74).

## Supporting information

Oberin and Petautschnig etal Supplementary Tables

## Declarations Acknowledgments

We thank Prof. Marnie Blewitt for critical comments on the manuscript, Monash Animal Research Platform staff for assistance with mouse care, Monash Histology Platform for assistance with slide scanning and the Monash Micro Imaging Facility and MHTP Medical Genomics Facilities for assistance and technical advice.

## Ethics approvals

All animal work was undertaken in accordance with Monash University Animal Ethics Committee (AEC) approval.

## Availability of data and materials

All RNA sequencing data have been deposited to the Gene Expression Omnibus (GEO) and are publicly available with accession number GSE210398. All other information is available from the corresponding author.

## Funding

This work was supported by grants and research funding from:

National Health and Medical Research Project Grant GNT1144966 (PSW, DKG, MvdB, DLA) Hudson Institute of Medical Research (PSW)

Victorian Government’s Operational Infrastructure Support Program.

Australian Government Research Training Program Scholarship (EGJ, RO and SP)

## Author contributions

Conceived and/or designed experiments: RO, SP, TT, ZQ, EJ, HB, PSW Performed experiments and/or analysed data: RO, SP, TT, NY, TTT, DF, ZQ, EJ, HB, PSW Bioinformatic analyses and/or comparisons with published datasets: ZQ, RO, SP, DLA, PSW Resources and/or supervision PSW DKG, MvdB, DLA, NAS

Writing - original draft: RO, SP, PSW

Writing - review & editing: RO, SP, EJ, PSW, NAS. All authors critically read and approved the manuscript.

## Supplementary Figure Legends

**Supplementary Figure 1.**
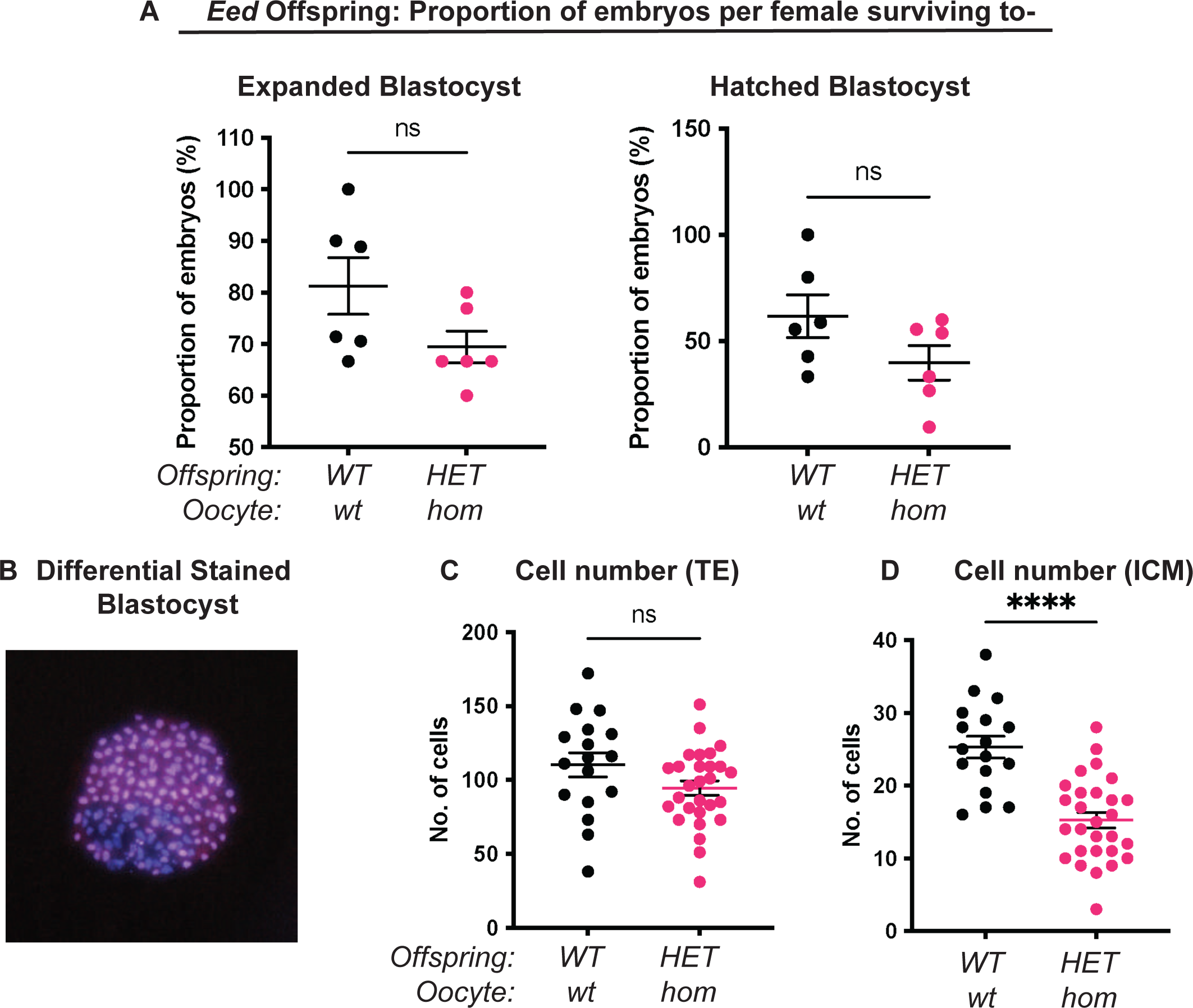
Maternal *Eed* deletion impacted pre-implantation development. (a) Proportion of embryos from *Eed-wt* or *Eed-hom* oocytes surviving to expanded blastocyst (left) and hatched blastocyst (right). Data represent the proportion of surviving embryos from each female, N = 6 females. **(b)** Example of a differentially stained blastocyst that has been treated with propidium iodide followed by bisbenzimide to identify purple cells representing inner cell mass (ICM) and pink cells representing trophectoderm (TE). **(c)** Number of TE cells per embryo. ns = no significant difference, P > 0.05, N = 17 embryos from *Eed-wt* oocytes, 28 embryos from *Eed-hom* oocytes. **(d)** Number of ICM cells per embryo. ****P < 0.0001, N = 17 embryos from *Eed-wt* oocytes, 28 embryos from *Eed-hom* oocytes. For **(a, c-d)** a two-tailed student’s t test was used, and error bars represent mean ± SD.

**Supplementary Figure 2.**
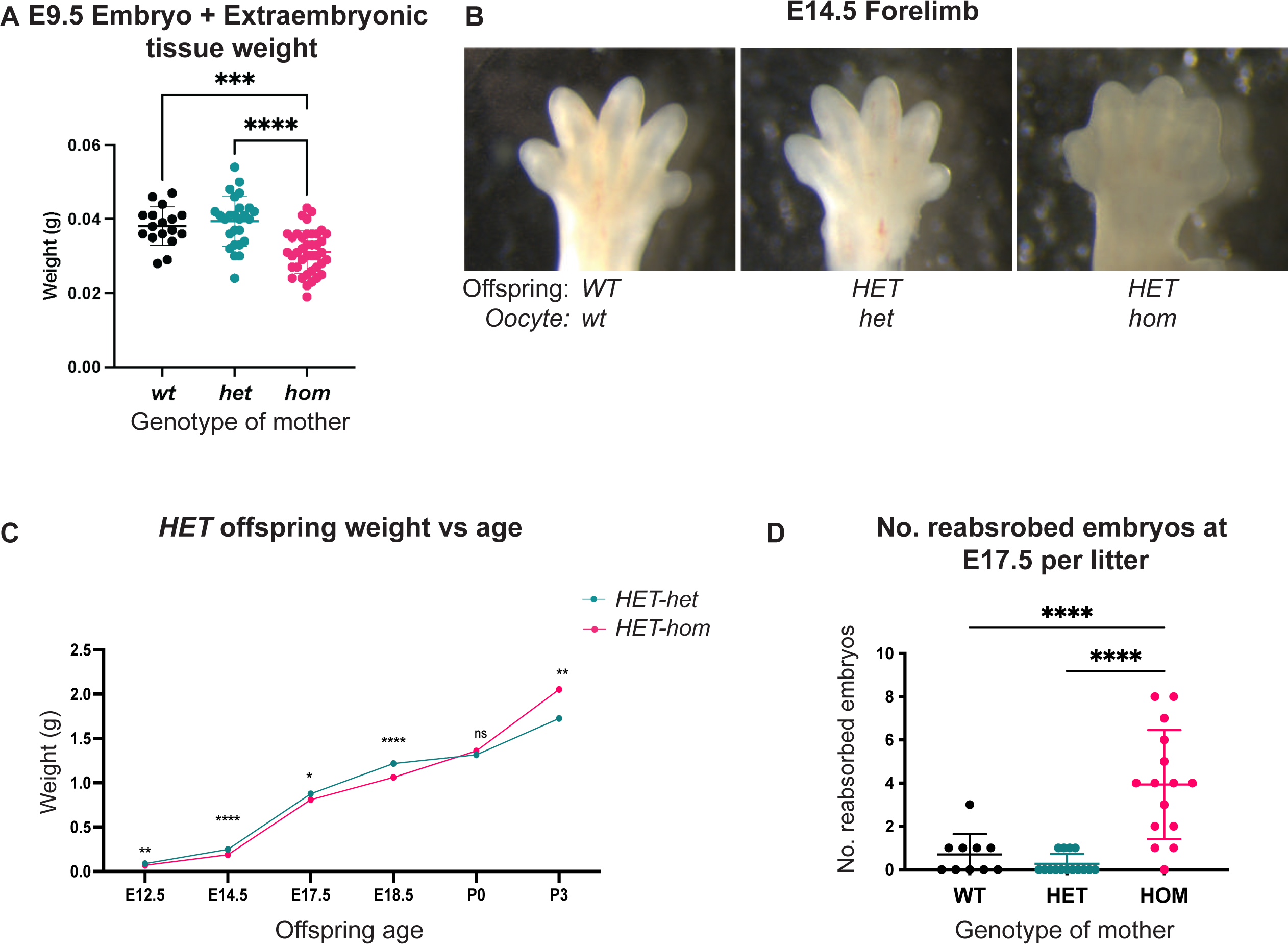
Loss of EED-dependent oocyte programming resulted in altered fetal and neonatal development. **(a)** Embryo and extraembryonic tissue weight at E9.5. ****P<0.0001, ***P<0.0005, N=17-40/genotype. **(b)** Representative wholemount images of E14.5 forelimbs revealing reduced inter-digital tissue regression in *HET-hom* offspring. **(c)** Isogenic heterozygous offspring average weight over time. *P<0.05, **P<0.005, ****P<0.0001 unpaired t-test, N=11-45/genotype. **(d)** Number of reabsorbed embryos per litter at E17.5. ****P<0.0001 N=10-15 litters/genotype. **(a, c)** statistically analysed using one-way ANOVA plus Tukey’s multiple comparisons.

**Supplementary Figure 3.**
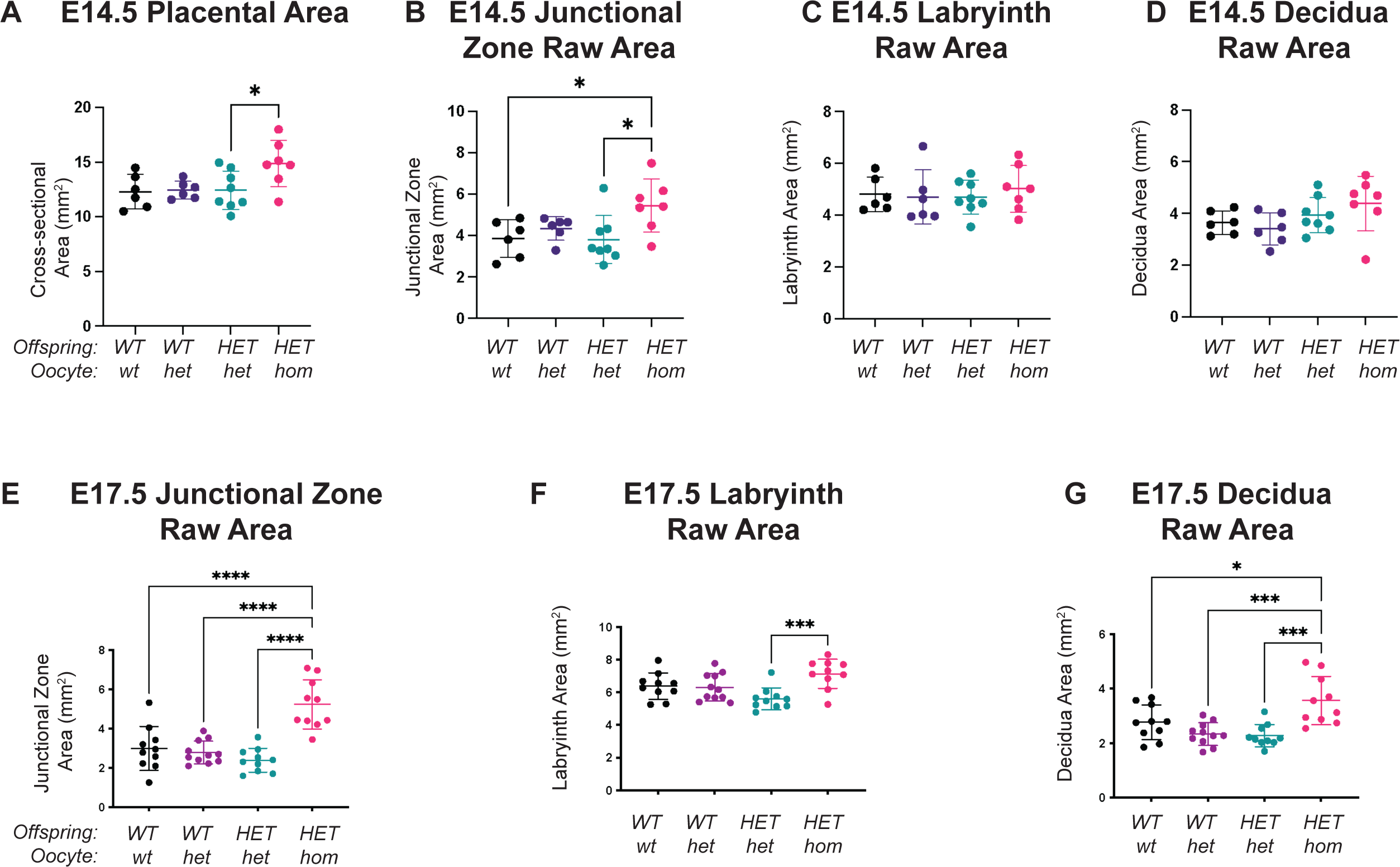
Complete deletion of *Eed* in the oocyte resulted in altered placental size. (a) E14.5 placenta cross-sectional area. *P<0.05, N=10/genotype. **(b-d)** Total area of E14.5 junctional zone, labyrinth and decidua. *P<0.05, N=10/genotype. **(e-g)** Total Raw area of E17.5 junctional zone, labyrinth and decidua. ****P<0.0001, ***P<0.0005, *P<0.05, N=10/genotype. Statistically analysed using one-way ANOVA plus Tukey’s multiple comparisons.

**Supplementary Figure 4.**
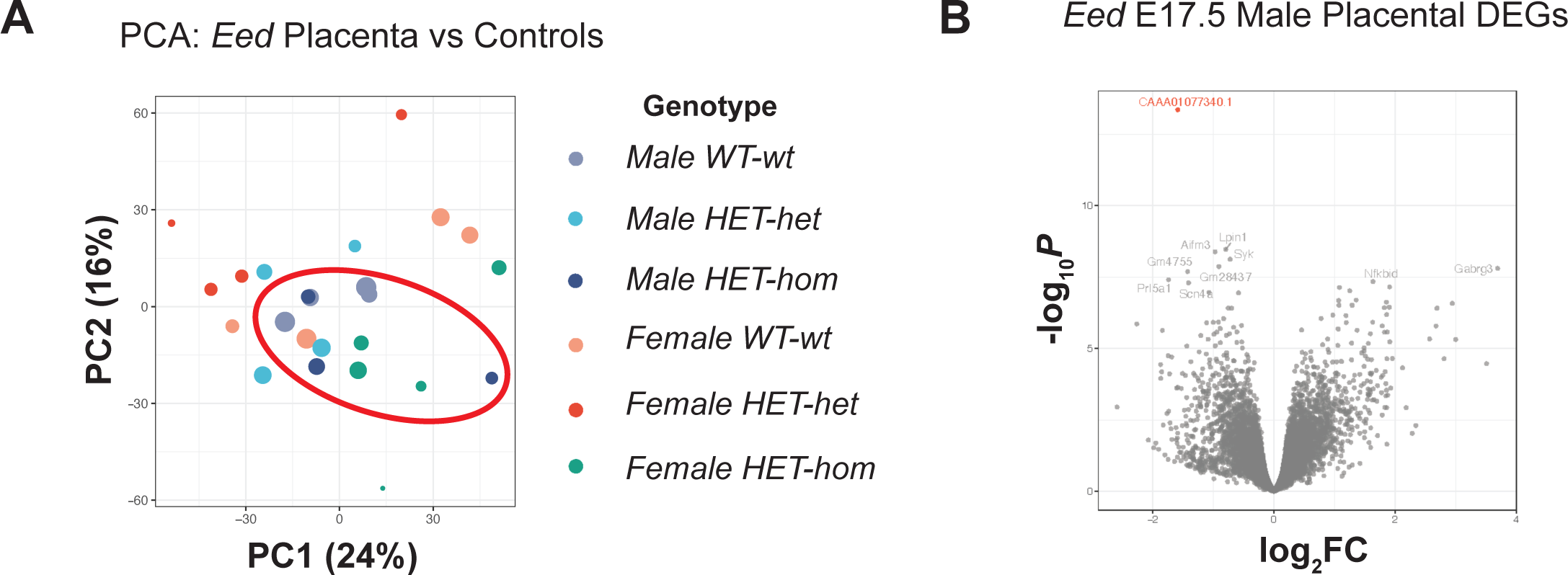
Loss of EED-dependent oocyte programming altered transcription in male *HET-hom* placenta. (a) Principal Component Analysis (PCA) for *HET*-*hom*, *HET-het* and *WT-wt* placenta. N=3-4/genotype. **(b)** Differential gene expression analysis of isogenic *HET-het* vs *HET-hom* placenta represented by a MD plot showing log2FC against statistical significance. Gene with FDR-adjusted FDR<0.05 is coloured in red. Deletion of *Eed* in the oocyte resulted in one DEG in male *HET-hom* placenta.

**Supplementary Figure 5.**
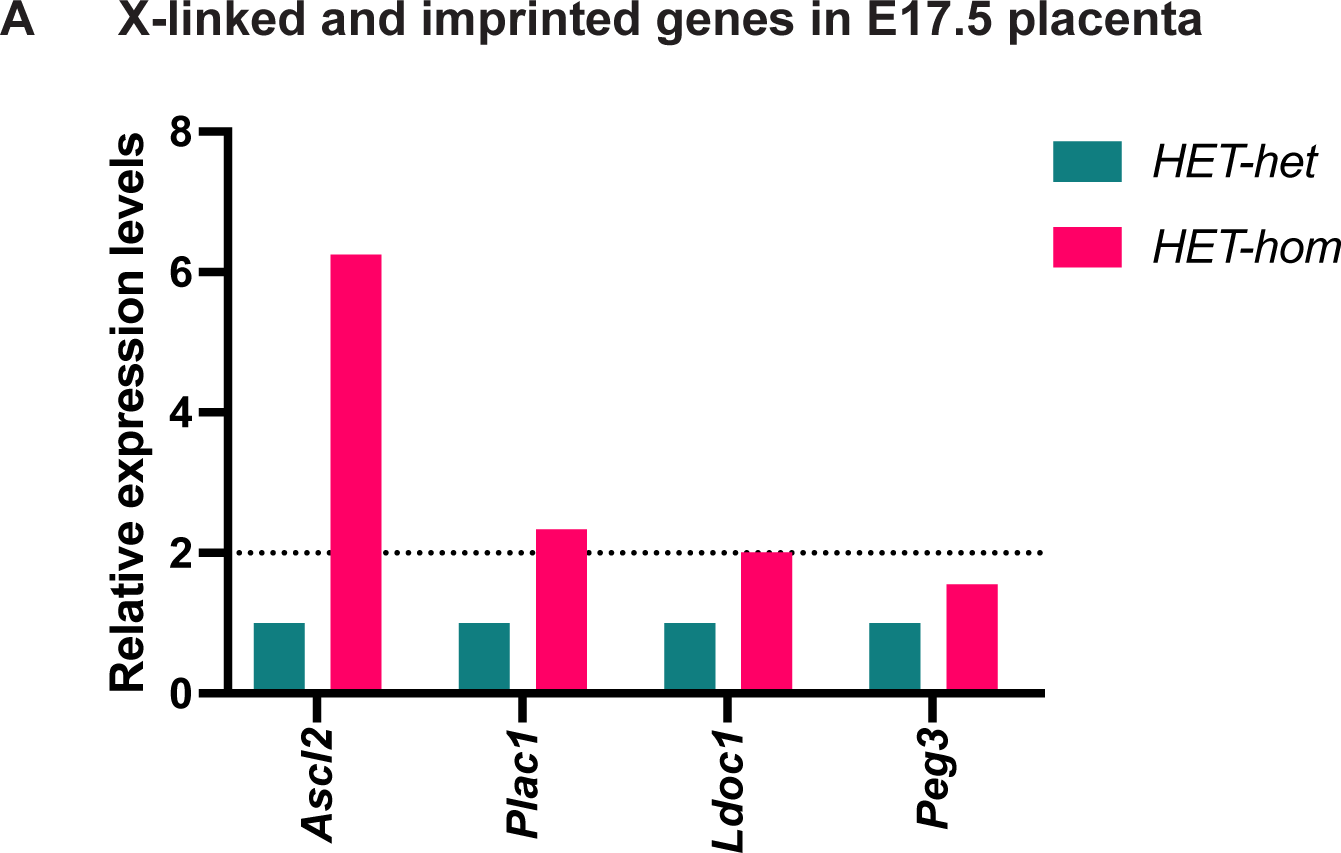
Loss of EED in the oocyte results in the de-repression of X-linked and imprinted genes that are important for placental development. (a) Relative expression levels of X-linked and imprinted genes in female E17.5 *HET-hom* placenta compared to female *HET-het* control (FDR<0.05).

**Supplementary Figure 6.**
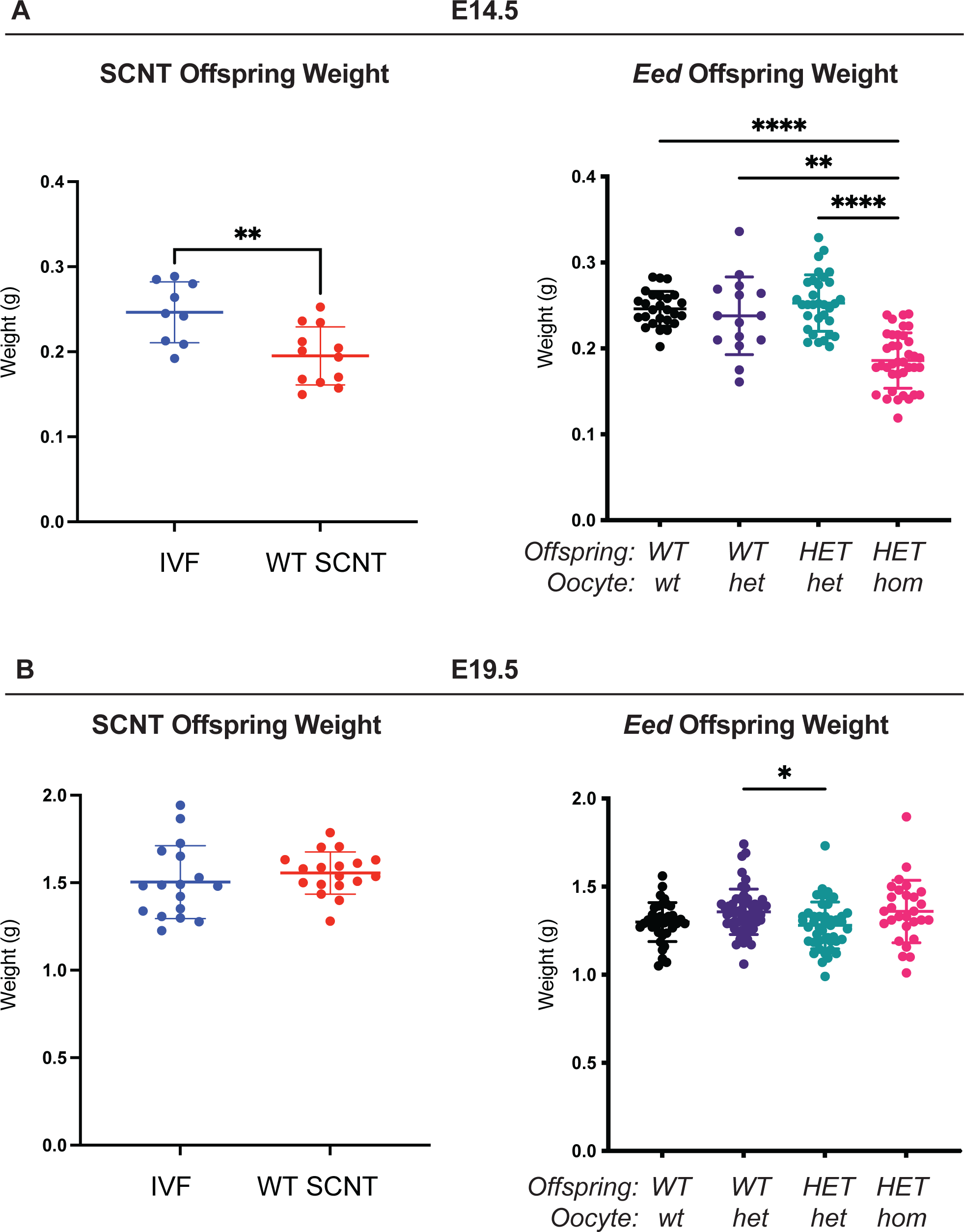
*HET-hom* offspring demonstrate similar growth profiles to SCNT mice. Comparison of body weights of SCNT mice from Xie, Zhang (27) and offspring from the *Eed-ZP3-*Cre mouse model at **(a)** E14.5 and **(b)** E19.5. ****P<0.0001, **P<0.005 and *P<0.05. Statistically analysed using one-way ANOVA plus Tukey’s multiple comparisons.

